# An ex vivo tissue model of cartilage degradation suggests that cartilage state can be determined from secreted key protein patterns

**DOI:** 10.1101/541375

**Authors:** Michael Neidlin, Efthymia Chantzi, George Macheras, Mats G Gustafsson, Leonidas G Alexopoulos

**Affiliations:** Department of Mechanical Engineering, National Technical University of Athens, Athens, Greece; Department of Medical Sciences, Uppsala University, Uppsala, Sweden; 4th Orthopaedic Department, KAT Hospital, Athens, Greece

**Keywords:** Osteoarthritis, cartilage degradation, multiplex protein profiling, proteomics, ex vivo model, cytokine interactions

## Abstract

The pathophysiology of osteoarthritis (OA) involves dysregulation of anabolic and catabolic processes associated with a broad panel of cytokines and other secreted proteins and ultimately lead to cartilage degradation. An increased understanding about the interactions of these proteins by means of systematic in vitro analyses may give new ideas regarding pharmaceutical candidates for treatment of OA and related cartilage degradation.

Therefore, first an ex vivo tissue model of cartilage degradation was established by culturing full thickness tissue explants with bacterial collagenase II. Then responses of healthy and degrading cartilage were analyzed by measuring protein abundance in tissue supernatant with a 26-multiplex protein profiling assay, after exposing them to a panel of 55 protein stimulations present in synovial joints of OA patients. Multivariate data analysis including exhaustive pairwise variable subset selection was used to identify the most outstanding changes in the measured protein secretions. This revealed that the MMP9 response is outstandingly low in degraded compared to healthy cartilage and that there are several protein pairs like IFNG and MMP9 that can be used for successful discrimination between degraded and healthy samples.

Taken together, the results show that the characteristic changes in protein responses discovered seem promising for accurate detection/diagnosis of degrading cartilage in general and OA in particular. More generally the employed ex vivo tissue model seems promising for drug discovery and development projects related to cartilage degradation, for example when trying to uncover the unknown interactions between secreted proteins in healthy and degraded tissues.

## Introduction

Osteoarthritis (OA) is a progressive disease involving mechanical, biochemical and genetic factors that disturb associations between chondrocytes and extracellular matrix (ECM), alter cellular metabolic responses and result in degradation of articular cartilage^1^. Prominent proteins associated with the pathophysiology of OA are pro-inflam-matory cytokines including the interleukins IL1a/b, IL6, IL8 and the tumor necrosis factor TNFa^1^. Anti-inflammatory cytokines such as IL4, IL10 and IL13 are also elevated in OA tissues^2^. Moreover, aggrecanases and matrix metalloproteinases (MMPs) that degrade the ECM as well as growth factor families of bone morphogenetic proteins (BMPs), fibroblast growth factors (FGFs) and transforming growth factors (TGFs) are all present in synovial joints of OA patients^3,4^. The fact that proteins with opposing effects are found in OA joints simultaneously (e.g. pro-inflammatory and anti-inflam-matory cytokines or matrix degrading enzymes and chondrogenic cytokines) suggests nontrivial inherent interactions between these proteins.

Targeting these players separately in order to reverse or suppress OA has been tested in many clinical studies in the past with very limited success. Monoclonal antibodies such as Adalimumab (anti-TNFa) and Canakinumab (anti-ILb), MMP inhibitors, and growth factor stimulators like Sprifermin (rhFGF18) have not been able to provide significant improvements until now or are still in clinical trials^5^. Such rather disappointing results support the hypothesis that targeting a single protein is not sufficient for a successful therapy. Therefore, a deeper understanding of the cytokine interaction network might be necessary to leverage drug discovery in OA.

One approach to achieve this aim is the use of antibody-based multiplexing assays that simultaneously measure the abundance of a broad panel of proteins in a biological sample. Applications of such assays related to OA and cartilage include re-constructions of chondrocyte cell signaling pathways based on phosphoproteomics and cytokine release data from 2D chondrocyte cultures^6^, cytokine releases after anabolic stimulations of 3D chondrocyte scaffolds^7^ and measurements of joint pathology dependent cytokine profiles in synovial fluids and cartilage tissues^8^. Notably, careful multivariate analyses of such protein measurements also have great potential as a tool for discovery of novel diagnostic biomarkers in the form of characteristic changes across subsets of the proteins studied. However, in order to enable systematic large-scale measurements of these protein secretion patterns, a sufficiently simple and cheap ex vivo tissue model of cartilage degradation (CD) is needed.

Many in vitro models of OA and CD have been developed in the past^9^. These differ in the tissue types used (monolayer cell cultures, 3D cell cultures or tissue explants) and the method chosen for OA induction (mechanical damage or chemical stimulation with pro-inflammatory cytokines). Some in vitro models use co-culturing with synovium, subchondral bone or other OA related tissues to represent more physiological conditions. Chemical induction of OA in an explant model often uses IL1b and/or TNFa to suppress the synthesis of proteoglycans and increase the release of MMPs that consequently cleave the collagen links of the ECM^9^. Another approach for modeling of CD associated with OA, recently proposed by Grenier et al.^10^, is pre-treatment of cartilage tissue explants by collagenase type II. Using this approach, cleavage of collagen II is directly induced and the ECM gets degraded, together with associated changes in surface morphology, decreases of tissue sulfated glycosoaminoglycan (s-GAG) content and a deterioration of the mechanical properties such as increased permeability and decreased Young’s moduli. As stated by the Grenier et al.^10^, this suggests that enzy-matic degradation with collagenase II can be used to simulate characteristic changes observed in early-stage OA.

Similarly to Grenier et al.^10^ we therefore used pure collagenase II pre-treatment as a degradation inducer in order to create a simplified ex vivo tissue model of CD. Using only collagenase II results in an oversimplification of the physiological conditions, but one can be sure that degradation will be achieved after a rather short treatment period. This model was established and then used in our protein profiling approach to understand how different protein stimuli affect the degraded state and to search for potential diagnostic biomarkers that can discriminate between healthy and degraded cartilage tissue. More specifically, we expanded the work by Grenier et al.^10^ by looking at protein secretion patterns after stimulation with major OA related cytokines/proteins, and evaluated the possibility to use these response patterns for determination of the cartilage state. As demonstrated below, this novel systematic approach revealed biomarkers with potential to be used for accurate detection/diagnostics of degrading cartilage. More generally, this approach was found to have potential to help uncovering the interactions of CD related proteins, and thereby also help accelerating drug discovery and development activities associated with OA..

## Materials and Methods Explant tissue model

### Model overview and workflow

The main idea of the ex vivo tissue model is to perturb healthy and degrading cartilage tissue with a set of OA related stimuli followed by a measurement of the tissue responses in terms of protein secretions. The resulting dataset is analyzed in order to compare the two different tissue states and pinpoint individual stimuli yielding different protein responses. Our hypothesis is that these protein responses depend on the tissue state (healthy or degrading) and thus can be used to distinguish between them.

Figure 1 illustrates the combined experimental-computational procedure with the individual steps described in more detail below.

**Figure 1:**
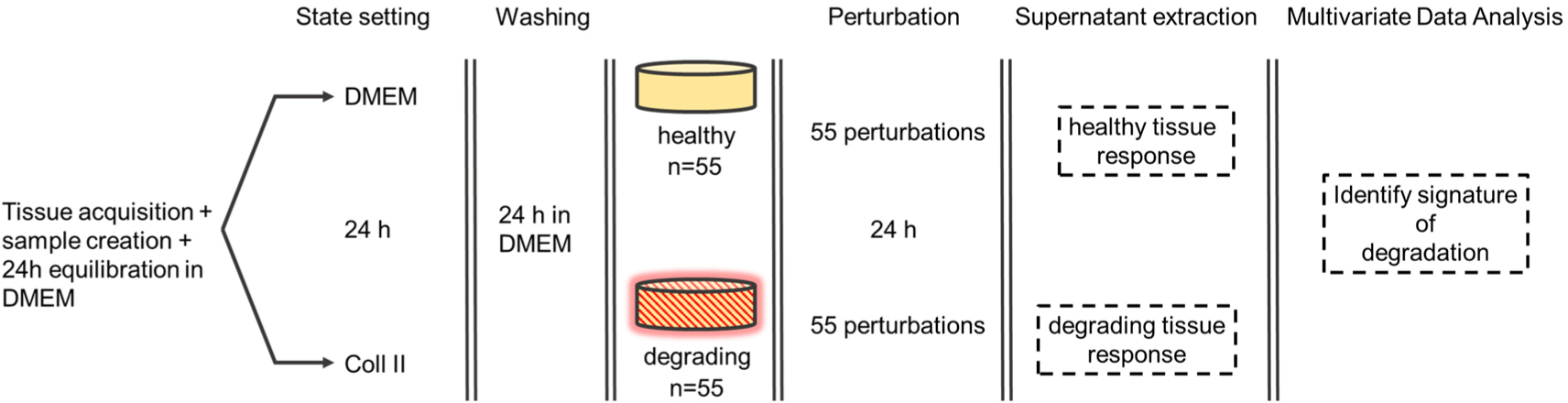
Overview of the novel combined ex vivo and in silico procedure introduced here. Cartilage tissue explants were retrieved from femoral heads of hip fracture patients and equilibrated for 24h in DMEM*. Tissue discs were pre-treated for 24h either with DMEM* or collagenase type II, and washed with DMEM* for 24h. Then the discs were stimulated for another 24h with 55 perturbations consisting of single proteins and pairwise combinations of some of them. Healthy and degrading responses were characterized in terms of multivariate patterns of secreted proteins measured in the supernatant after the stimulation.

### Explant isolation, state setting and washing

Cartilage tissue samples were obtained from the femoral heads of patients (n=5, age 72-94, 1 male and 4 females) undergoing total hip replacement due to fracture (n=3, P1-P3) or OA (n=2, P4 and P5) with patient’s informed consent and protocol approved by the responsible ethics committee. For an overview of how the samples were used in the different experiments performed, see Table 1. The samples were examined macroscopically to determine the locations of intact cartilage. Cartilage parts showing signs of degradation were excluded. Most of the cartilage from OA patients was acquired from the lateral part of the central superior femoral head. In the fracture cohort, almost all parts of the femur head could be used. Femur heads were rinsed with PBS, cartilage without subchondral bone was removed and placed into high glucose DMEM (Dulbec-co’s Modified Eagle Medium) supplemented with 10% FBS, 1% Penicillin/Streptomy-cin and 1% fungizone (BioCell Technology LLC, Irvine, CA), denoted by DMEM*. Cartilage disc samples of 3mm diameter were created with a biopsy punch and let to equilibrate in DMEM* for 24h. Then the tissue samples were placed in either fresh DMEM* or DMEM* with collagenase type II, activity 125 units/mg, (MP Biomedicals, Santa Ana, CA) of 2 mg/ml for 24h. To see the effect of different collagenase concentrations on the cytokine/protein secretion of the cartilage explants, concentrations of 1 and 4 mg/ml were also used. The three collagenase concentrations were applied in duplicate. Finally, before starting the perturbations of the resulting healthy and degrading cartilage samples, a washing step of 24h in fresh DMEM* media was included.

**Table 1:** Overview of the experiments and replicate number per tissue state for all experiments performed in the study. For details, see the main text.

| Experiment and number of replicates | Patient | Origin | Result location |
| --- | --- | --- | --- |
| Preliminary analysis:<br>Steady state (n=2 per tissue state) | P2 | fracture | Supplementary<br>Material 2 |
| Preliminary analysis:<br>Batch effect and CV (n=1 per patient) | P1 + P2 | fracture | Supplementary<br>Material 3 |
| Protein release after 3 different colla-<br>genase II concentrations (n=2 per group ) | P2 | fracture | Main text |
| Healthy and degrading tissue response<br>(n=1 per tissue state) | P2 + P3 | fracture | Main text |
| Histology (n=2 per tissue state) | P4 | OA | Supplementary<br>Material 5 |
| GAG release (n=3 per tissue state) | P4 | OA | Supplementary<br>Material 5 |
| Collagen II content (n=3 per tissue state) | P5 | OA | Supplementary<br>Material 5 |
| Mechanical test (n=3 per tissue state) | P5 | OA | Supplementary<br>Material 5 |

### Perturbation and supernatant extraction

A total number of 13 proteins for stimulation were selected as they have been reported being present in OA^1,2^. These included IL1a, IL1b, TNFa, IL6, IL8, IL4, IL10, IL13, BMP2, FGF2, IGF1, TGFb1 and MMP9 (PeproTech EC Ltd, London). These proteins were separated into the following two groups; [IL1a, IL1b, TNFa, IL6, IL8, MMP9] and [IL4, IL10, IL13, BMP2, FGF2, IGF1, TGFb1]. All possible single treatments and all possible pairwise combinations from these two groups were used as a perturbation set, resulting in 55 different stimuli. The concentrations used were chosen from prior experiments and already existing studies^6,7,9^. Detailed information on the experimental design can be found in the Supplementary Material 1. A stimulation time of 24h was chosen based on studies about the transient behavior of tissue responses, which can be found in Supplementary Material 2. In Supplementary Material 3, the results show that no strong batch effect due to patient-to-patient variability could be observed and also that the coefficient of variation of the assay was acceptable (<25%) for 24 out of 26 proteins. Cartilage discs were put in 96 well microplates and individually stimulated with 260 µl of media. After the stimulation 80µl of the supernatant was retrieved and cytokine releases were measured with the FlexMap 3D platform (Luminex Corp. USA). The supernatant of an unstimulated disc (cultured in DMEM* during the perturbation step) was taken as control. Blank measurements to evaluate the experimental noise for each protein were included as well. All steps were conducted in a humidified incubator at 37 ο C and 5% CO_2_. The rather high number of stimuli reduced the possibility of having biological replicates as 55 cartilage explants were needed for one run of single measurements. As our main aim was to discover and compare response patterns of cartilage explants and not to uncover new biological mechanisms, we decided to accept the drawback of having a low number of replicates with simultaneously having a broad panel of stimuli. Thus, single measurements were collected after the stimulations of 55 untreated cartilage discs of patient P2 and 55 collagenase II (2 mg/ml) treated cartilage discs of patient P3.

## Experimental techniques

### Multiplex ELISA

The Luminex xMAP technology is an antibody-based suspension array technology measuring protein abundance in a sample for a set of predetermined proteins. Detailed background information can be found in the review of Alexopoulos et al.^11^ A library of 26 protein releases (PEDF, CXCL11, IL13, ZG16, IL4, GROA, IFNG, CYTC, IL8, IL17F, IL12A, TNFa, IL1a, TFF3, ICAM1, IL10, FST, S100A6, CXCL10, PROK1, CCL5, IL20, TNFSF12, BMP2, FGF2, MMP9) was measured in the supernatant.

### Histology evaluation

Two DMEM* and two collagenase II (2 mg/ml) treated cartilage discs were taken after the washing step (Figure 1), fixated in PBS with 10% formalin, decalcified and embedded in OCT. Histological evaluation with toluidine blue staining following standard protocols was performed^12^.

### Mechanical testing

In order to evaluate the change of mechanical properties of collagenase II (2 mg/ml) treated samples, three DMEM* and three collagenase II (2mg/ml) treated tissue samples were taken after the washing step and tested with the Bose Electroforce 3100 (Bose, Framingham, MA). Stress-relaxation tests and measurement of the equilibrium stress were used to obtain information about the compression stiffness of the material^13^. Initially, samples were pre-loaded with a force of F=0.1N. Then an instantaneous ramp displacement of 5% of the initial height was applied and the relaxation of the force over time was measured until a dynamical equilibrium was reached. The procedure was repeated for a total of three loading steps. Equilibrium stress was calculated as the engineering stress σ=F/A_0_ with A_0_≈7.07 mm^2.

### GAG release

The extracellular release of sulfated glycosaminoglycans was measured spectrophotometrically via a Dimethylmethylene Blue (DMMB) assay^14^ using the Varioscan LUX multimode microplate reader (Thermofisher Scientific Inc., USA). As s-GAGs belong to the main constituents of the ECM^10^ an increased presence in the explant supernatant can be directly related to increased ECM destruction. The GAG release was quantified for two groups. One group was treated with DMEM* and the other group was treated with collagenase II (2 mg/ml). 50 µl of the supernatant was extracted at t=2h, 6h, and 24h. The measured s-GAG concentration was normalized to tissue wet weight. n=3 discs were chosen per group.

### Collagen II content

Collagen II is the main constituent of the ECM^10^, thus reduction can be directly related to ECM destruction. Tissue collagen II content was quantified as shown previouly^15^ with a hydroxyproline assay kit (Abcam, Cambridge, UK). One group was treated with DMEM* for 24h and the other group was treated with collagenase II (2 mg/ml) for 24h. n=3 discs were chosen per group and tissue collagen content was normalized to tissue wet weight.

## Data post-processing, analysis and statistical tests

The multiplex ELISA experiments delivered median raw fluorescence intensities (MFI) for each marker in the cytokine release dataset resulting in (55 perturbation +1 control)*26 (proteins) =1456 data points for the healthy and degrading tissue, respectively. Additionally duplicate blank measurements were included for each protein. MFI values of measurements below the average of the blanks were deleted and imputed based on a nearest neighbor algorithm^16^ as implemented in the function `knnimpute` provided by Matlab (MathWorks, Natick, USA). In addition, the same imputation algorithm was used to replace the saturated MFI values obtained for the particular protein(s) used for stimulation. For example, when IL1a is used for stimulation, then the corresponding MFI value is saturated and thus replaced via imputation.

Meaningful multivariate data analyses required normalized MFIs that allow comparisons between plates and between cytokines/proteins. The normalized difference *D(i,j,p)* for stimulus i and cytokine/protein j present on plate p was determined as:

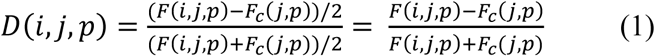

where F(i,j,p) denotes the MFI from cytokine j for stimulus i on plate p and F_c_(j,p) denotes the signal from the untreated control well for cytokine j on plate p. The normalized values are all restricted to the interval [−1,+1] where the value +1 is obtained when F(i,j,p) ≫F_c_(j,p), the value -1 when F(i,j,p) ≪F_c_(j,p) and the value 0 when F(i,j,p)=F_c_(j,p). Matlab and R^17^ were used for the post processing and the analysis of the data. Principal Component Analysis^18^ (PCA) and k-means clustering^19^ were used to explore the data. Subsequently, supervised methods in the form of optimal orthogonal system discriminant analysis (OOS-DA)^20^ and an exhaustive pairwise variable subset selection procedure were employed in order to identify the most discriminative pairs of proteins.

### OOS-DA, optimal orthonormal system for discriminant analysis

OOS-DA^20^ may be viewed as a multi-output generalization of Fisher’s linear discriminant analysis^21^ where multiple orthogonal linear projections are created sequentially, each maximizing Fisher’s separation criterion between the classes of interest. OOS-DA has been used previously to design a new type of classification method^22^ and for batch correction in mass spectrometry based metabolomics^23^. The OOS-DA was performed by means of Matlab code developed in-house.

### Exhaustive pairwise variable subset selection

The between-class separation s_b_ and the within-class separation s_w_ needed to calculate the separation score J_separation_ = s_b_/s_w_ used to find the most discriminative pairs of measured proteins are defined in equations 2 and 3.

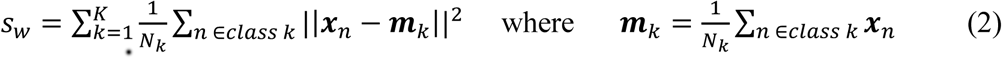

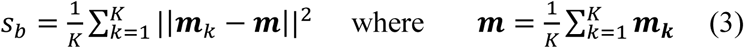

In the analysis performed here we used K=2, corresponding to healthy and degraded samples, respectively. **x**_n_ denotes the n^th^ sample (column vector) and N_k_ denotes the number of samples belonging to class k. In order to reduce the risk of overfitting, the actual score used to identify the most discriminating protein pair was calculated as the maximum across 100 values of J_separation_ obtained using a resampling approach where each value was obtained by using a stimuli subset consisting of 80% of the stimuli in the original dataset.

### Statistical comparisons of individual treatments

Statistical comparisons between individual treatments were done with the unpaired two-sided t-test at significance level p=0.05.

Table 1 provides an overview of the sample and replicate number per tissue state for all experiments performed in the study. Furthermore the tissue source (patients P1-P5), its origin (fracture or OA), and the location of the associated results presented in this work are listed.

## Results

### Collagenase II treatment induces protein releases observed in clinical OA

In order to investigate the individual protein secretions of collagenase treated tissue, raw MFIs were compared between untreated (DMEM*) and treated (Col) samples. The MFIs obtained from the three concentrations were pooled together. Cohen’s d, defined as d=(m_1_-m_2_)/s where m_i_ are the class means and s is the pooled standard deviation of the dataset, was taken as the measurement of the effect size. The identified cytokines are presented in Table 2, sorted by decreasing effect size. Out of the 11 proteins with significant differences between control and collagenase treated samples, 4 (bold, Table 2) were found in many OA related studies. These were TNFa and IFNG that are related to inflammation and innate immunity responses as well as the two anti-inflammatory cytokines IL4 and IL13^24^. CXCL11^25^, IL17F^26^, and TNFSF12^27^ (TWEAK) are also considered contributors of OA. The remaining 4 cytokines TFF3^28^, PEDF^29^, CCL5^30^ and S100A6^31^ can be considered underreported players. However every protein has been shown to play a role in at least one study. In summary, these results suggest that cartilage discs treated with collagenase increase the secretion of particular proteins that also have been observed to have increased levels in clinical OA studies

**Table 2:** Protein secretions showing significant differences between DMEM* (n=3) and collagenase (n=6) treated samples. Sorted according the effect size measured in terms of Cohen’s d.

| Protein | Function/Involvement | p | Cohen's d |
| --- | --- | --- | --- |
| **IL4** | anti-inflammatory | <0.001 | 5.7 |
| TFF3 | unsure | <0.001 | 5.2 |
| **IFNG** | innate immunity | 0.018 | 4.2 |
| PEDF | multiple functions | 0.001 | -3.2 |
| **TNFα** | pro-inflammatory | 0.001 | 3.1 |
| CCL5 | immunity response | 0.007 | 3.0 |
| **IL13** | anti-inflammatory | 0.002 | 2.6 |
| S100A6 | Ca2+ binding | 0.009 | 2.4 |
| CXCL11 | T cell chemoattraction | 0.011 | 2.3 |
| IL17F | pro-inflammatory | 0.012 | 1.4 |
| TNFSF12 | pro-inflammatory | 0.03 | 1.3 |

### Tissue state can be determined based on measured protein patterns

PCA of healthy and degrading tissue responses with subsequent k-means clustering (k=2) based on the first 4 PCA dimensions (82% of variance covered) was performed. The responses plotted in the resulting 2D space of the first two PCs are presented in Figure 2.

**Figure2:**
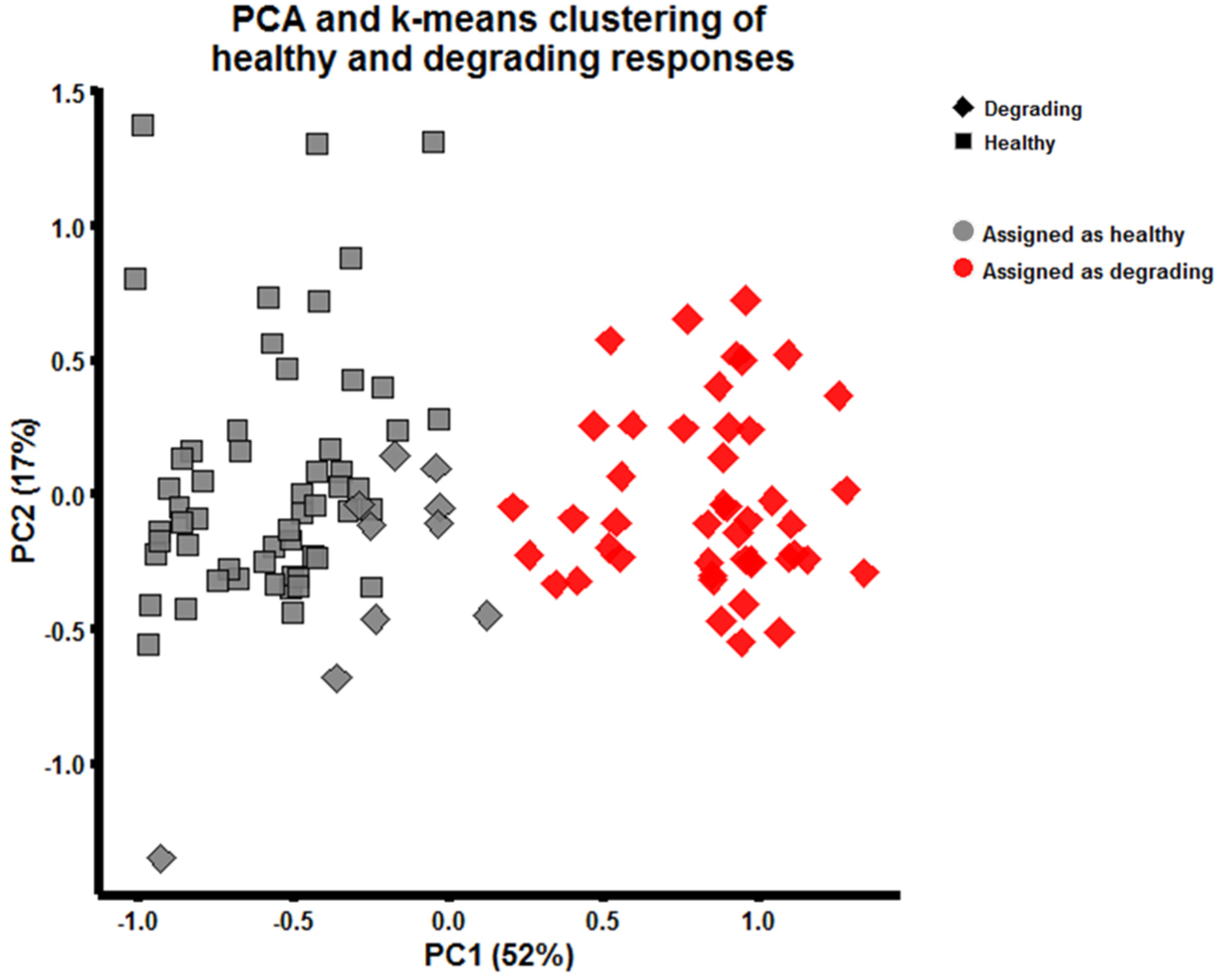
PCA score plot (first two principal components covering 52% and 17% of the variance, respectively) of normalized MFIs. The shape of a marker indicates its true tissue (sample) group. The colors reflect the unsupervised category assignments made by k-means clustering (k=2) category assignments. As the samples assigned to the gray color mainly belong to the healthy group, it has been given the label “assigned as healthy” while the red samples has been given the label “assigned as degrading”

The shapes of the markers in Figure 2 represent the true tissue (sample) groups. There are 55 squares (healthy perturbed discs) and 55 diamonds (degrading perturbed discs) in the PC1-PC2 plane. The two colors gray and red represent the two categories identified by the k-means algorithm (k=2). The subsequent manual assignment of them as “assigned as healthy” and “assigned as degrading” protein secretion patterns was based on the fact that the majority of the members of the gray cluster belongs to responses from the healthy tissue and the majority of the red cluster belongs to responses the degrading tissue.

Figure 2 also shows that protein secretions after perturbations lead to the formation of two distinct clusters in the PC1-PC2 plane, indicating that tissue state can be determined based on measured protein patterns. In particular, the coordinate along the PC1-axis seems suitable for classification of the cytokine responses as “healthy” or “degrading”. This suggests that it is interesting to look at the elements (loadings) of the corresponding eigenvector in order to determine which of the measured protein changes are most useful for separation between healthy and degrading tissue. The individual loadings of each protein to PC1 are shown in Supplementary Material 4, Figure S1. Increases of MMP9 and FGF2 as well as decreases of CXCL10, ZG16 and FST would cause a shift along PC1 from degrading to healthy responses.

### Levels of IFNG and MMP9 are promising for tissue state discrimination

PCA is an unsupervised method that disregards any class information. Therefore it is not guaranteed that the resulting linear projection provides an optimal solution in terms of discrimination between healthy and degrading tissues. Notably, 10 out of the 55 degrading tissue responses get assigned to the wrong cluster (Figure 2, gray diamonds). Therefore, supervised discriminant analysis in the form of OOS-DA^20^ was used instead, to identify the proteins most accountable for the changes between degrading and healthy responses. The full analysis can be found in Supplementary Material 4. Briefly, applying OOS-DA identified that there were many linear combinations of the 26 measured protein changes that lead to better separation than using the corresponding PCA projections (Supplementary Material 4, Figure S2). Contrary to the analysis of the PC1 loadings, no outstanding influence of FGF2 on class separation could be observed using OOS-DA. Therefore, instead of performing dimensionality reduction using OSS-DA that yields ambiguous results, a straightforward variable subset selection procedure was employed in the form of an exhaustive search across all plausible subsets of two proteins where the optimal protein pair regarding class discrimination was identified. The resulting scatterplot of this pair (MMP9, IFNG) together with a linear decision boundary separating the two classes is shown in Figure 3. The arrow indicates the direction in which the degrading tissue responses should be moved in order to coincide with the healthy tissue responses.

**Figure3:**
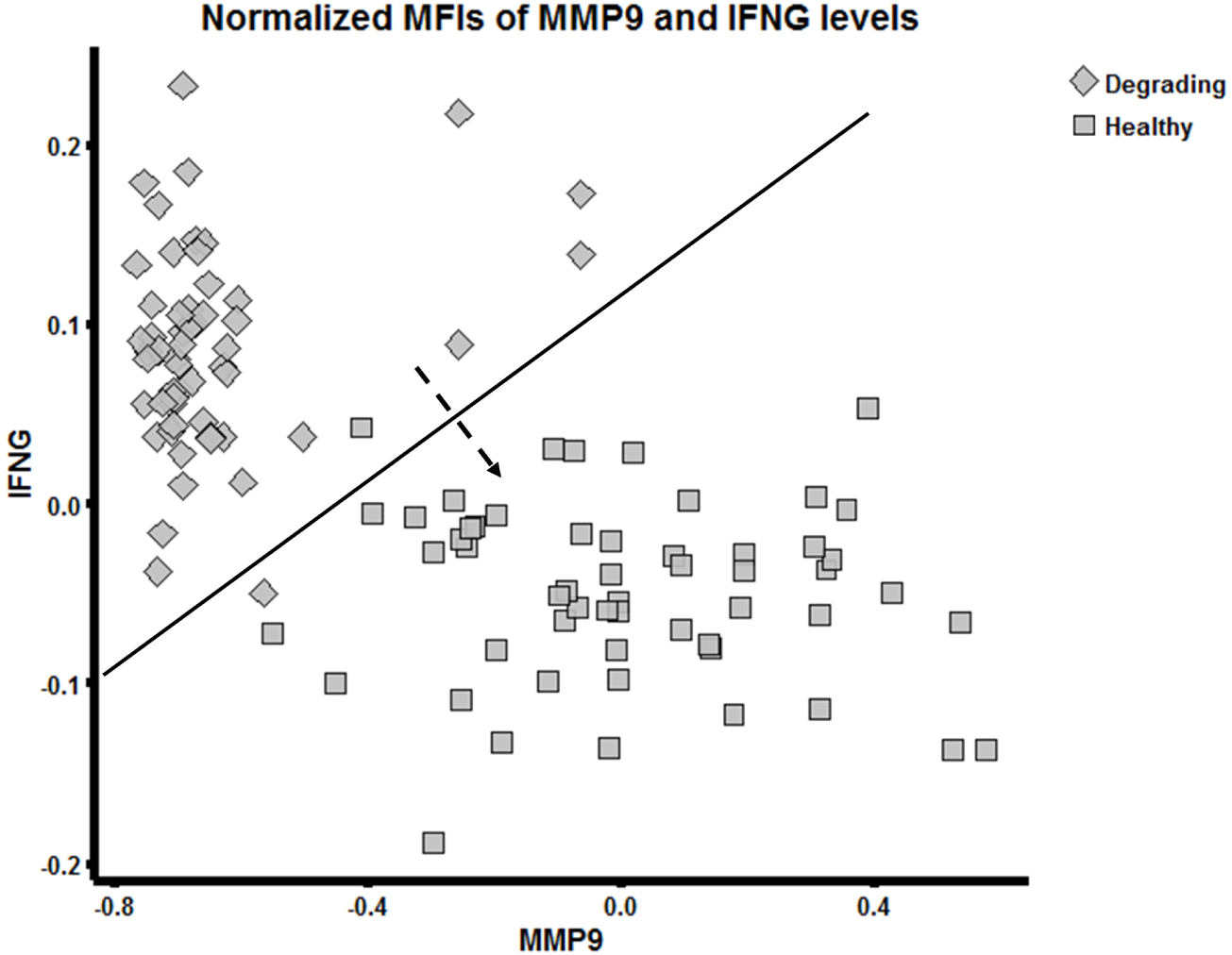
A scatterplot of IFNG and MMP9 levels (normalized MFIs) of healthy and degrading responses. The line indicates the decision boundary between healthy and degrading tissue. The arrow indicates the desired direction that the degrading responses should move to.

In summary stimulation of healthy and degrading tissue with a set of 55 stimuli lead to changed secretion levels of various proteins, which were evaluated to discriminate between the two tissue states using different approaches. An exhaustive search across all pairwise combinations of protein levels identified MMP9 and IFNG, among several other pairs, as promising for class discrimination.

### Collagenase II treatment induces OA like phenotype changes

The results of histology evaluation, mechanical testing, as well as measurements of GAG release and collagen II content in seemingly healthy cartilage samples taken from OA patients are presented in Supplementary Material 5. These analyses show that collagenase II treatment leads to: (i) structural changes of cartilage (after 24h at concentration of 2 mg/ml) characterized by tissue fissuring and fibrillation, (ii) significant (p<0.001) decrease of tissue collagen II content, (iii) significant decrease (p<0.001) of the elastic modulus, (iv) significant (p<0.001) increase of GAG release into the supernatant. Taken together, these results partly confirm the results reported by Grenier et al.^10^, in general showing that collagenase II treatment induces OA like phenotype changes.

## Discussion

The main aim of this study was to measure and characterize cytokine/protein release patterns in an ex vivo model of CD, created by exposing healthy cartilage to the ECM degrading enzyme collagenase type II. First, it was shown that the induced ECM degradation resulted in secretions of proteins related to OA observed in various clinical studies (see Table 2). Secondly, the in vitro responses of healthy and degrading tissue to a broad set of OA related protein perturbations were investigated. Multiplexed ELISA measurements of 26 secreted proteins were subject to exploratory (PCA, k-means) multivariate data analysis. These analyses showed that it is possible to successfully distinguish between healthy and degrading samples for a majority of the samples (Figure 2). However, 10 out of 55 responses from the degrading tissue were assigned incorrectly and supervised OOS-DA showed that there are many other linear projections, other that those obtained using PCA (see Supplementary Material 4, Table S1), which give almost perfect separation for all samples. Finally, an exhaustive variable subset selection algorithm was employed to identify pairs of proteins that have the most promising discrimination between the classes. The changed levels of MMP9 and IFNG delivered the best distinction between the classes according to the separation criterion used (Figure 3). Notably, the cloud of squares in the scatter plot (Figure 3) may be interpreted as a characteristic distribution (“fingerprint”) showing how a healthy cartilage responds to a panel of 55 protein perturbations. Similarly, the cloud of diamonds in Figure 3 reflects how a degraded cartilage responds to the same perturbations. Ideally a successful drug treatment would make the degraded cartilage produce the same cloud as the healthy cartilage, meaning that the cloud of diamonds moves to sit on top of the cloud of squares, as indicated by the striped arrow in Figure 3.

In many studies on in vitro models for OA related drug treatment, the typical experimental readouts are glycosaminoglycan (GAG) and collagen II content. In addition prototypic biomarkers such as matrix metalloproteinases (MMPs) or inflammatory factors such as IL1a/b or TNFa are also often used^32^. Looking at such few readouts might be too simplistic given the outstanding complexity of the human biological systems. In particular, this might lead to misleading conclusions in general, for example overlooking truly efficient single drugs and drug combinations. Therefore, in our approach we measured the changed release of 26 proteins simultaneously. Then we used the collected data in order to characterize the molecular processes associated with CD, including how they can be used for diagnosis and/or to accelerate drug discovery and development.

Both our molecular and phenotypic results support the idea of Grenier et al.^10^ to use collagenase II to induce an “OA-like” state. For the phenotypic results, we used healthy looking parts of tissues obtained from OA patients and confirmed that collagenase II treatment results in expected/desired OA-related properties like structural changes of cartilage, tissue fissuring and fibrillation, decreased tissue collagen II content, decreased elastic modulus, and increased GAG release into the supernatant (see Supplementary Material 5). In their work, Grenier et al. performed collagenase II treatment for 45-120 min, and claimed it to induce similarities with early stages of OA^10^. In our work the collagenase II treatment was extended to cover 24h. The panel of released proteins after such a treatment in our study (Table 2) indicates similarities with late rather than early stages of OA, when comparing with clinical studies. However, this difference in interpretation may simply be due to the fact that there is yet no clearly defined distinction between early and late stages of OA.

### Limitations

The systems biology approach introduced here based on protein profiling of CD used 55 stimulations per cartilage condition (healthy and degraded). Since a patient donation usually results in less than 100 cartilage samples, the tissue responses recorded in this study did not come from the same donor. More specifically, healthy perturbed (gray squares in Figure 2) and degrading perturbed (red diamonds in Figure 2) came from two different patients (P2 and P3). Therefore, one cannot exclude that there is a batch effect that can explain some of the differences observed. However, as shown in Figure S1 of Supplementary Material 3, when looking at perturbed healthy cartilage from P1 and P2, no batch effect is observable. Future studies should reduce the number of perturbations employed in each experiment, thereby allowing for using cartilage samples from the same patient for both states (healthy and degrading) on the same 96-well plate. Additionally, using duplicate measurements on the same experimental plate would also be preferable. The relatively high number of 55 different perturbations was used to increase the probability to observe different protein releases from healthy and degrading tissue, whilst accepting a statistically weaker significance of the biological differences observed. Thus, the framework and results presented here should be considered as a proof-of-concept that will be followed by more optimized experiments in the future, which will be limited to a smaller set of stimulations.

A second limitation of the study reported here is that important OA related proteins such as disintegrin and metalloproteinases with thrombospondin motifs ADAMTS 4 and 5 and the matrix metalloproteinase MMP13 were not included. In the current study these proteins were not measured due to the lack of reliable assays and challenges in terms of cross-talk. Additionally, it would be interesting to quantify to which extent different stimuli affect the collagen II content of the explants.

Despite these limitations we believe that the novel systemic ex vivo and in silico approach introduced here presents a viable way to investigate compound treatments for CD in general, and related to OA in particular. Thus, we have shown that a more sophisticated systemic analysis at the molecular level is feasible, and that it provides a more detailed molecular picture of what happens during CD in terms of proteomic responses to cytokine/protein stimulations.

In this study we presented an ex vivo model of CD for systemic profiling of interactions between stimulating and responding proteins. Measurements of key protein patterns seem useful to make statements about the condition of an unclassified tissue sample. Future work will investigate changes in tissue responses after pre-treatments with chemical agents (drug candidates) that have the potential to inhibit ongoing CD.

## Acknowledgement

MN acknowledges financial support from the German Research Foundation (DFG) via the scholarship ‘‘Forschungsstipendium” (PN: 387071423). All authors have nothing to disclose.

Author contribution statement
MN and LA conceived and designed the study. Based on the patient samples provided by GM, MN performed all the wet lab experiments. MN also performed most of the final data analyses and data visualizations. EC and MG introduced and implemented, the MFI normalization procedure, the OSS-DA analysis, the exhaustive variable subset selection method as well as the idea to replace saturated protein readouts by means of imputed values. The interpretations of the resulting data were made jointly by all authors. LA supervised the project. MN drafted the manuscript based on inputs from all co-authors. All authors read and approved the final version of the manuscript.

