## Supplementary Information 1 for "An ex vivo tissue model of cartilage degradation suggests that cartilage state can be determined from secreted key protein patterns"

**Experimental layout**

The experimental layout for stimulation of tissue discs together with the applied concentrations is shown in Figure S1. The bold borders describe the plate, each cell contains a cartilage sample. 130 µl of the top row and the left column were used in order to create 55 (+1 control) stimuli of final concentrations half of the shown values.

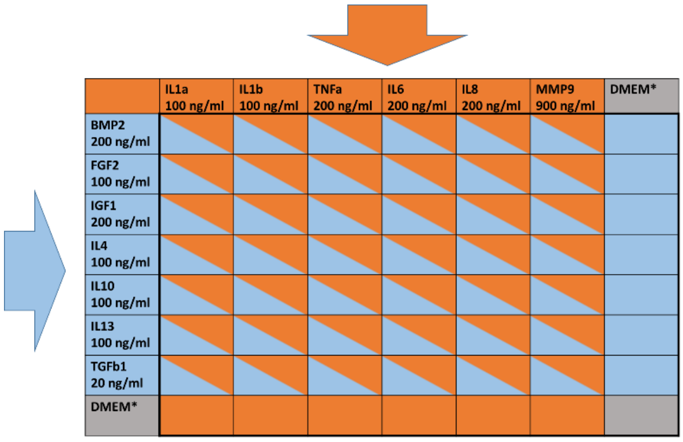


Figure S1: Layout for tissue stimulation. Top row and left column are mixed with an equal amount of 130 µl in order to produce 55 different stimuli.

The stimuli were chosen because of their major and well-known involvement in osteoarthritis. Table 1 describes their function with the respective reference.

| Name | Full name and function | Ref. |
|---|---|---|
| IL1a | major pro-inflammatory cytokine | [1] |
| IL1b | major pro-inflammatory cytokine | [1] |
| TNFa | major pro-inflammatory cytokine | [1] |
| IL6 | dual role/regulatory: pro-inflammatory cytokine and anti-inflammatory potential | [1] |
| IL8 | dual role/regulatory: pro-inflammatory cytokine and anti-inflammatory potential | [1] |
| MMP9 | gelatinase, degrades extracellular matrix | [2] |
| BMP2 | chondrogenic - major role in development of bone and cartilage | [1] |
| FGF2 | growth factor – supports tissue growth in many tissues | [1] |
| IGF1 | growth factor - supports tissue growth in many tissues | [1] |
| IL4 | major anti-inflammatory cytokine | [1] |
| Il10 | major anti-inflammatory cytokine | [1] |
| IL13 | major anti-inflammatory cytokine | [1] |
| TGFb1 | major growth factor | [1] |

[1] Mary B Goldring. Osteoarthritis and cartilage: the role of cytokines. *Current rheumatology reports*, 2(6):459–465, 2000.
[2] Brandon J Rose and David L Kooyman. A tale of two joints: the role of matrix metalloproteases in cartilage biology. *Disease markers*, 2016, 2016.
