## Supplementary Information 2 for "An ex vivo tissue model of cartilage degradation suggests that cartilage state can be determined from secreted key protein patterns"

In order to select an appropriate stimulation time an experiment investigating the time-dependent behavior of tissue responses was conducted. Stable cartilage discs were cultured with Col II [2 mg/ml], IL1a, IL1b, IL8 [all 50 ng/ml] and DMEM* then supernatant was retrieved at t=6h, 24h, 48h and 72h.

Multiplex ELISA with a 26-plex as stated in the main manuscript was used to measure the protein abundance.

To consider the relative changes, the normalized MFI values were calculated according the following equation:

$$D_{t_{n+1}-t_n}(i,j) = \frac{F(i,j,t_{n+1}) - F(i,j,t_n)}{F(i,j,t_{n+1}) + F(i,j,t_n)}$$

where F denotes the raw MFI values for stimulus *i* and cytokine *j* and *n* stands for the respective time points 6,24,48 and 72h. Thus, relative changes between 24h vs. 6h, 48h vs. 24h and 72h vs. 48h were calculated for 4 stimuli (+1 DMEM) and 26 cytokines. Hierarchical clustering with row and column reordering was applied, see Figure S1:

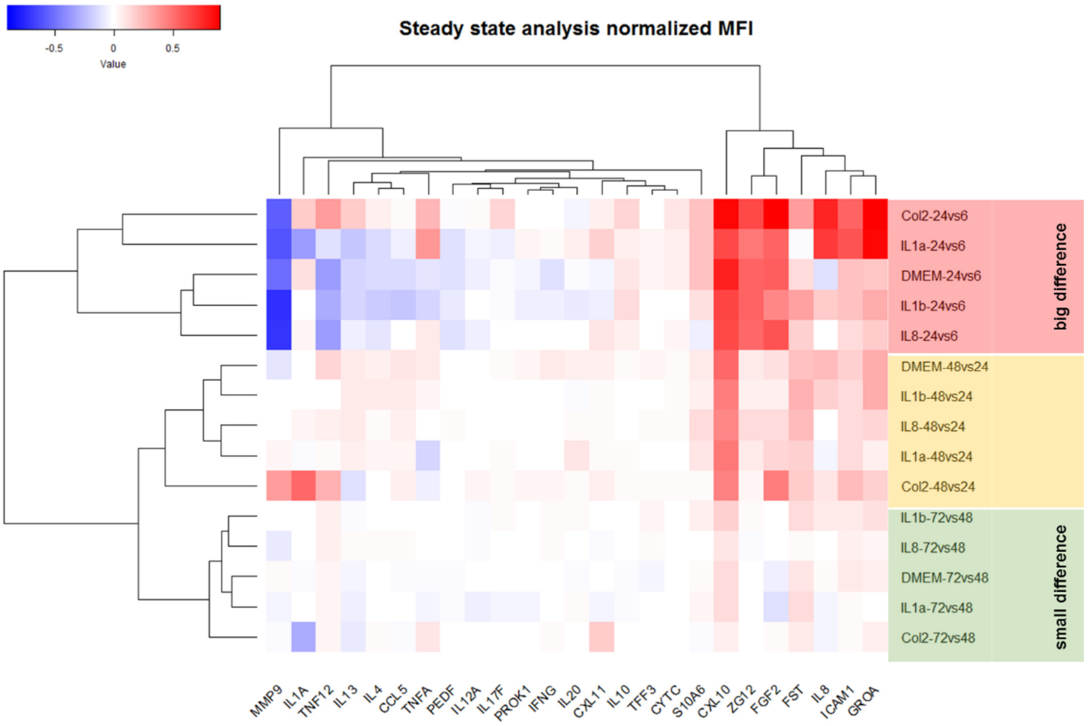


Figure S1: Hierarchical clustering of normalized MFIs for steady state analyses. White color corresponds to no changes for the respective stimulus and cytokine and the time point pair. Blue and red correspond to reductions and increases of MFI values.

As the first observation it can be seen that stimuli (rows) of the corresponding groups (24vs6, 48vs24 and 72vs48) cluster together without any outliers. Thus, the induced cytokine releases are consistent throughout the stimuli for all time points. The 24h vs.

6h group is located in the big difference cluster. This means that there are big changes between the protein releases after 6h and after 24h of stimulation and the system has not reached its steady state after 6h of stimulation. On the contrary, the 72h vs. 48h cluster exhibits almost no changes (white color in heatmap) between the two time points. The 48h vs. 24h group shows increases in CXCL10 for all stimuli and increases in MMP9, IL1A and TNFSF12 for the collagenase II stimulation.

Considering the fact that there are not many changes observed between 48h and 24h of the duration of 24h was chosen.
