## Supplementary Information 3 for "An ex vivo tissue model of cartilage degradation suggests that cartilage state can be determined from secreted key protein patterns"

### Analysis of patient-to-patient and assay variation

In order to confirm that the results reported in the main text are not easily explained by patient specific batch effects and/or by very noisy and stimulation specific measurements, the following analyses of patient-to-patient and assay variation were performed. At first, it was assessed how much patient-to-patient variability affects the overall results by comparing the 55 different 26-dimensional MFI profiles collected after stimulating 55 healthy cartilages from two different patients. Additionally the coefficient of variation (CV) for the 26-plex assay used in the experiments was computed in order to see the assay variation per protein and if this variation is different between untreated and treated samples..

#### Patient-to-patient variability

Healthy cartilage discs of P1 and P2 were perturbed with 55stimuli+1control with the experimental setting mentioned in Supplementary Material 1 for 24h. 80µl of the supernatant was retrieved and cytokine (set of 26 proteins as mentioned in the main text) releases were measured with the FlexMap 3D platform. Then PCA was performed on the entire dataset and the results were plotted in the resulting PC1-PC2 plane (Figure S1).

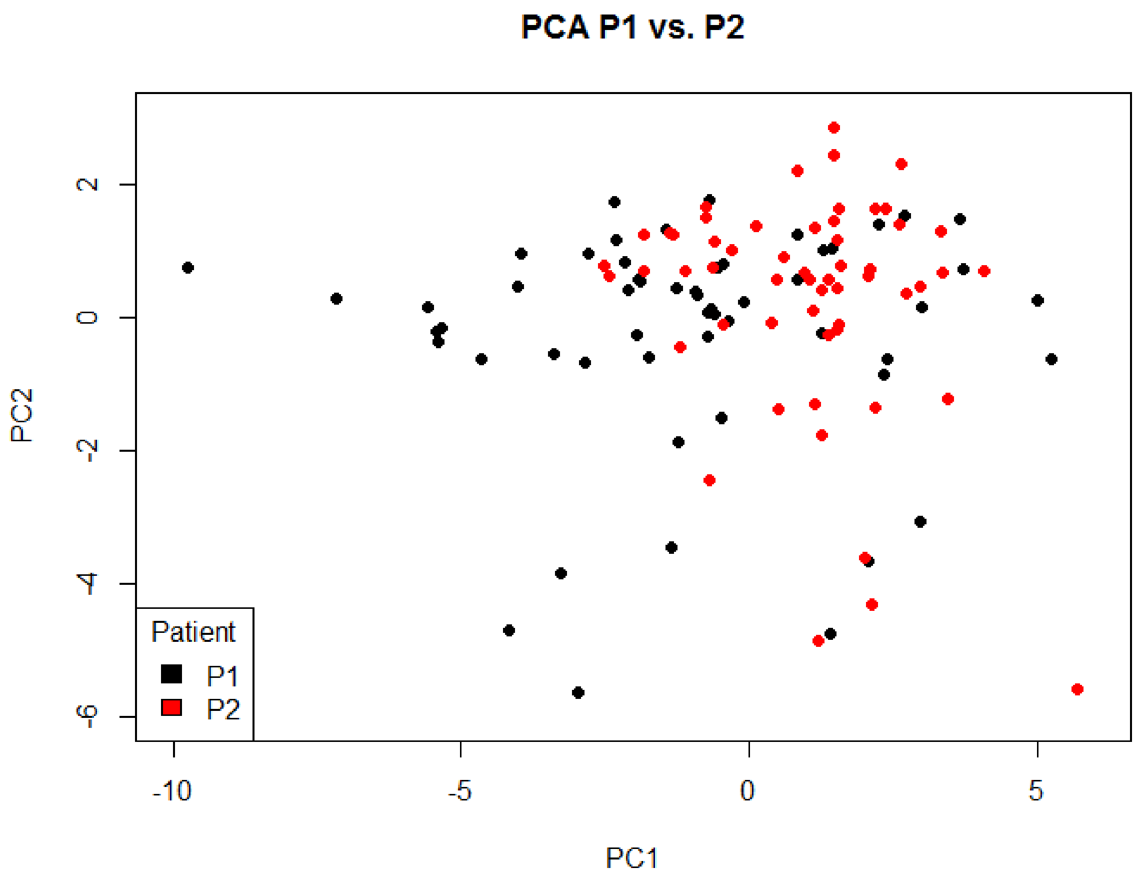


**Figure S1:**Score plot after PCA of cytokine releases of patients 1 (black) and 2 (red). As there is no apparent cluster formation in this plot, there are no signs of batch (patient specific) effects.

Figure S1 shows that no distinct clustering is observable in the two datasets of P1 and P2. In other words, there are no signs of batch (patient specific) effects. .

**Analysis of the assay coefficient of variation**

The quality of the 26-plex assay was assessed by looking at the coefficient of variation (CV) per protein in one stimulated and one unstimulated case. First, supernatant from one cartilage disc (patient P1) treated for 24h with IL-1α (stimulated case) was extracted. Next the supernatant was diluted (3:1) to enable triplicate 26-plex measurements during one experimental run. Further on, supernatant from one DMEM* (unstimulated) treated cartilage disc was extracted, diluted (3:1) and protein abundance was measured. A library of 26 protein releases (PEDF, CXCL11, IL-13, ZG16, IL-4, GROA, IFN-γ, CYTC, IL-8, IL-17F, IL-12A, TNF-α, IL-1α, TFF3, ICAM1, IL-10, FST, S100A6, CXCL10, PROK1, CCL5, IL-20, TNFSF12, BMP-2, FGF-2, MMP-9) was measured. In the stimulated case (IL-1α treatment) the measurements of IL-1α (26-plex) described fully saturated conditions and were disregarded. Thus, supernatant from one cartilage disc treated for 24h with IL-1β was also included and the results of IL-1α measurements were considered. The choice of IL-1α and IL-1β was based on the fact that these stimuli produced many responses of the cartilage tissue in terms of protein releases into the supernatant. The results of the CV analysis are found in Figure S2.

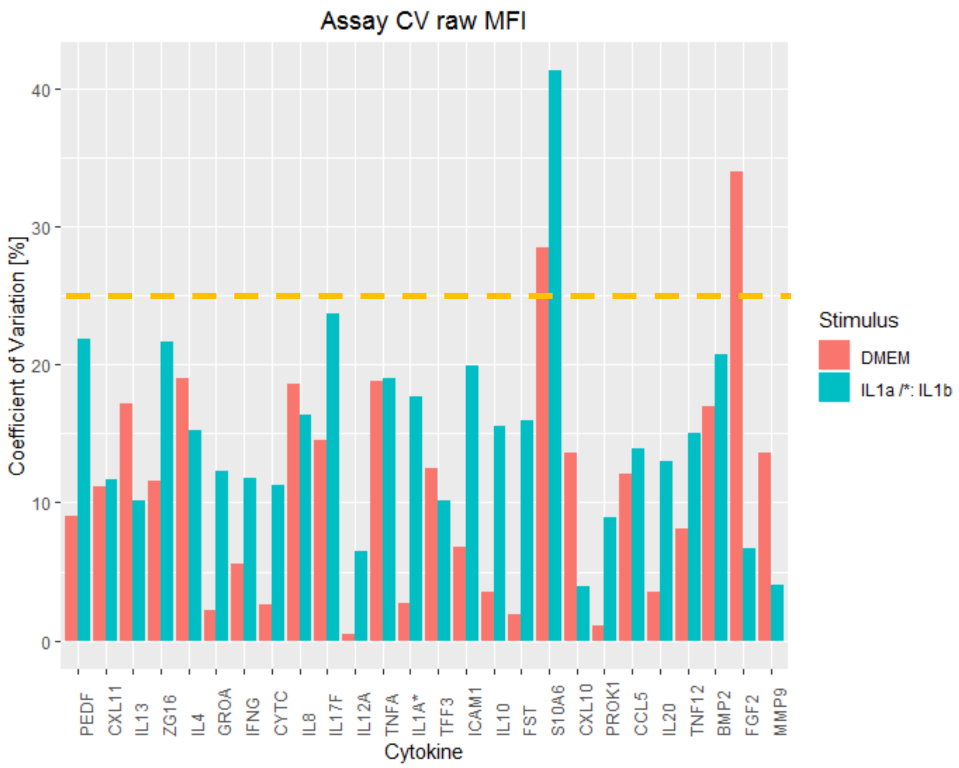


Figure S2: Coefficient of variation (CV) of raw MFI values in the cases of DMEM* and IL-1α /IL-1β . Yellow dashed line represents the threshold of 25%. Experiments were run in triplicate. No apparent difference in variability between untreated (DMEM*) and treated cartilage is observed and all CVs are below 25% except for FGF-2 and S10A6.

In 2 out of 26 cases the CV is above 25%, this is observed for S10A6 (stands for protein S100A6) and FGF-2. However MFI values of the FGF-2 measurements were ~200 for the stimulated and unstimulated cases, meaning being in the region of experimental noise. For all other measurements DMEM* showed a lower CV, which is a comforting observation as the normalization procedure relates all measurements to the control well (DMEM* treatment), see equation 1 in the main text.

In summary, the CV analysis revealed that measurements of S10A6 and FGF-2 are unreliable as their CV values are above 25%. The remaining proteins of the assay produce more reliable measurements (below CV 25%). Moreover, as the CV values are roughly the same for the DMEM* and IL-1α /IL-1β based stimulations, the CV analysis also shows that there is no apparent treatment specific effect on the assay variability. As a limitation one has to keep in mind that IL-1α / IL-1β treatment did not lead to a release of all 26 proteins. However it is not possible to find a stimulus that leads to a release of all proteins with a reasonable time and cost frame.
