## Supplementary Information 4 for "An ex vivo tissue model of cartilage degradation suggests that cartilage state can be determined from secreted key protein patterns"

Additional figures supporting the results section of the main manuscript are shown below.

**Healthy and degrading tissue response – Loadings of principal component 1**

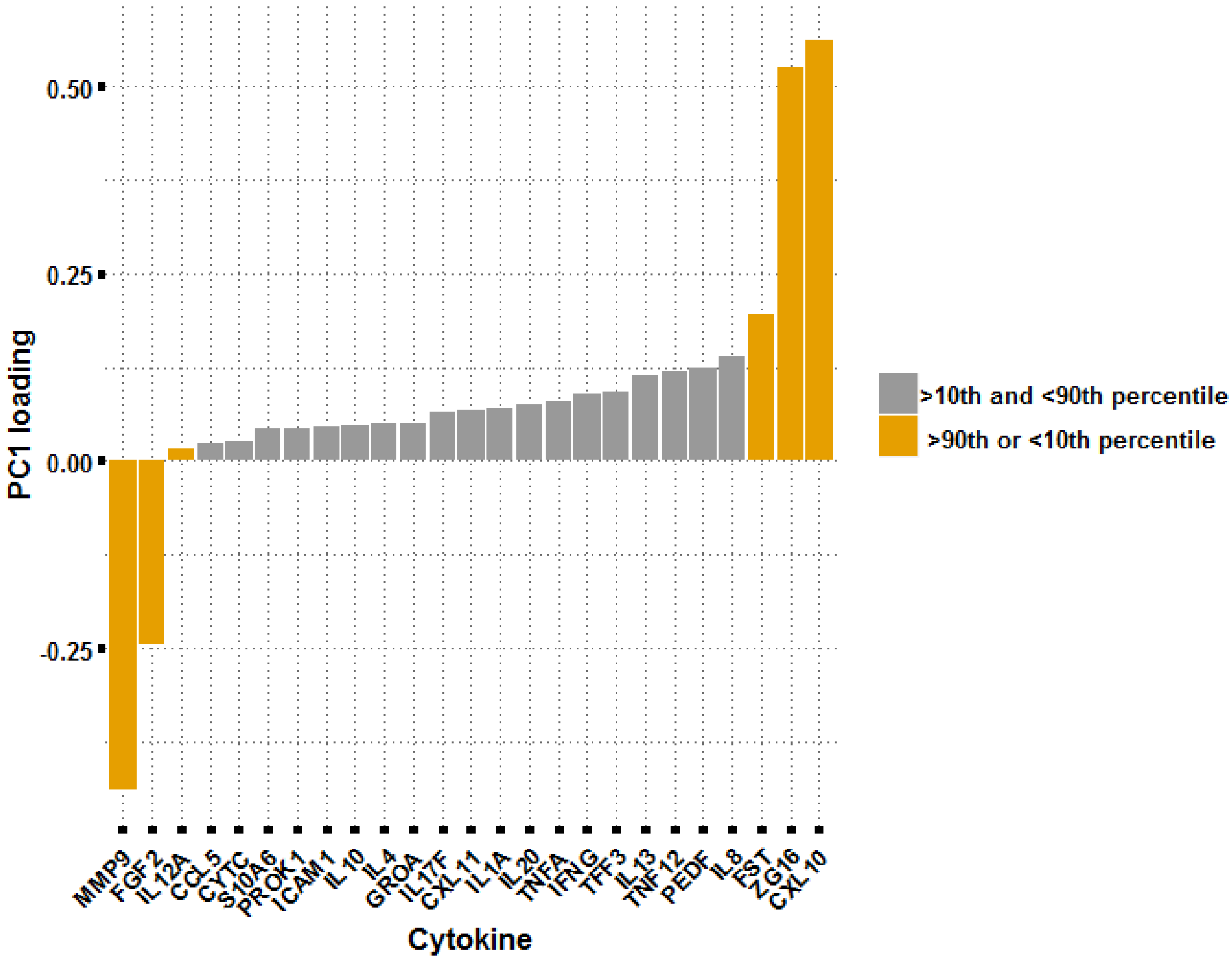


Figure S1: Individual loadings in PC1. The values below the 10th percentile and above the 90th percentile are highlighted in yellow.

### Optimal orthonormal system for discriminant analysis

*Complete separation is possible using many linear projections*

The results from PCA (Figure 2 in main manuscript) and loading analysis shown in Figure S suggest that certain changes in protein levels after stimulation are more informative than others for discrimination between degraded and stable cells. However, employing instead supervised optimal orthonormal system for discriminant analysis (OOS-DA) reveals that there are numerous linear projections (and linear combinations of them) that lead to complete separation between the collected stable and degenerated cytokine profiles, see Figure S (left) for one example. The corresponding projection vectors are shown in Figure S (right). For details regarding OOS-DA, see Materials and methods of the main text. This great variety of linear projections offering perfect class separation means that one cannot with great certainty determine the importance of individual proteins only by looking at the magnitude of the corresponding weights (loadings). In other words, ranking the proteins according to the magnitudes of their corresponding weights (loading) may be confusing and/or misleading. In order to avoid this potential pitfall, instead of analyzing weights (loadings) of linear projections, we performed an exhaustive search across all plausible subsets of 2 proteins to determine the pairs being the best for separating the two classes.

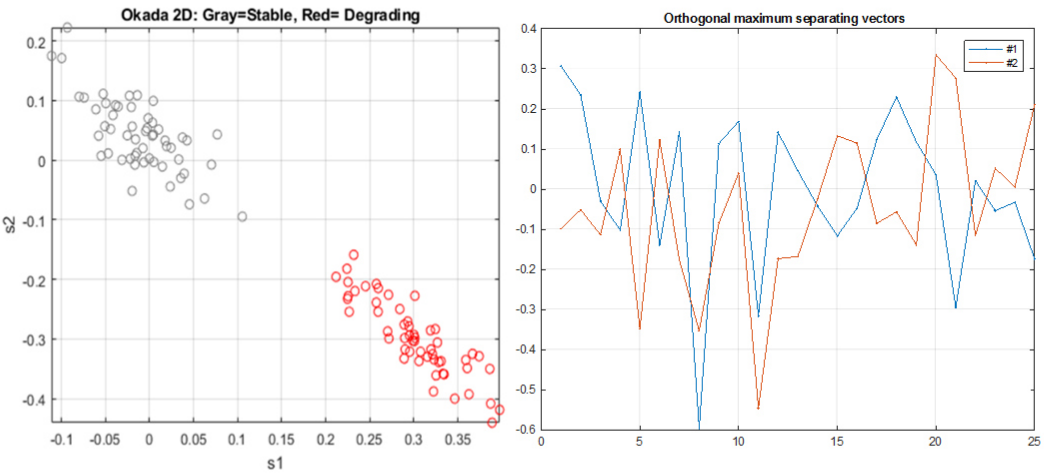


Figure S2: *Left*: Linear compression down to two dimensions using OOS-DA showing complete separation of the classes and therefore better results compared to the corresponding compression in Figure 2 (main manuscript) obtained using PCA. *Right*: Projection vectors obtained from OOS-DA used to obtain the total separation shown in the scatterplot presented in the left subfigure.

*Finding the most discriminative pairs of measured proteins*

To find the most discriminative pairs of measured proteins via an exhaustive search, the separation score

$$J_{separation} = s_b/s_w$$

was used where $s_b$ is a measure of the spread between the two classes (stable and degraded) and $s_w$ is a measure of the spread within the two classes for details see Materials and Methods. Histograms and scatterplots corresponding to selected single proteins and pairs identified using this approach are presented in Figure S (top) together with a matrix indicating the separation scores for all pairs (bottom). Most importantly, this shows that MMP9 alone is outstanding for discrimination between the classes. Moreover, the three first subfigures in the upper part show that one would like to find drugs that simultaneously increase MMP9 and decrease IFNG responses in degraded cells. From the fourth (bottom right) subfigure one finds that it would be great if the drugs also simultaneously decrease both IFNG and CXCL11 responses in degraded cells.

Considering the observations from Figure S it becomes obvious that there is a big amount of pairs that provide good separation (everything with $J > 0.6$).

Table 1 lists all the pairwise combinations with $J_{separation} > 0.6$ sorted by decreasing score. It is obvious (as from Figure 3) that almost all combinations including MMP9 and many combinations including CXCL10 produce satisfying class discrimination.

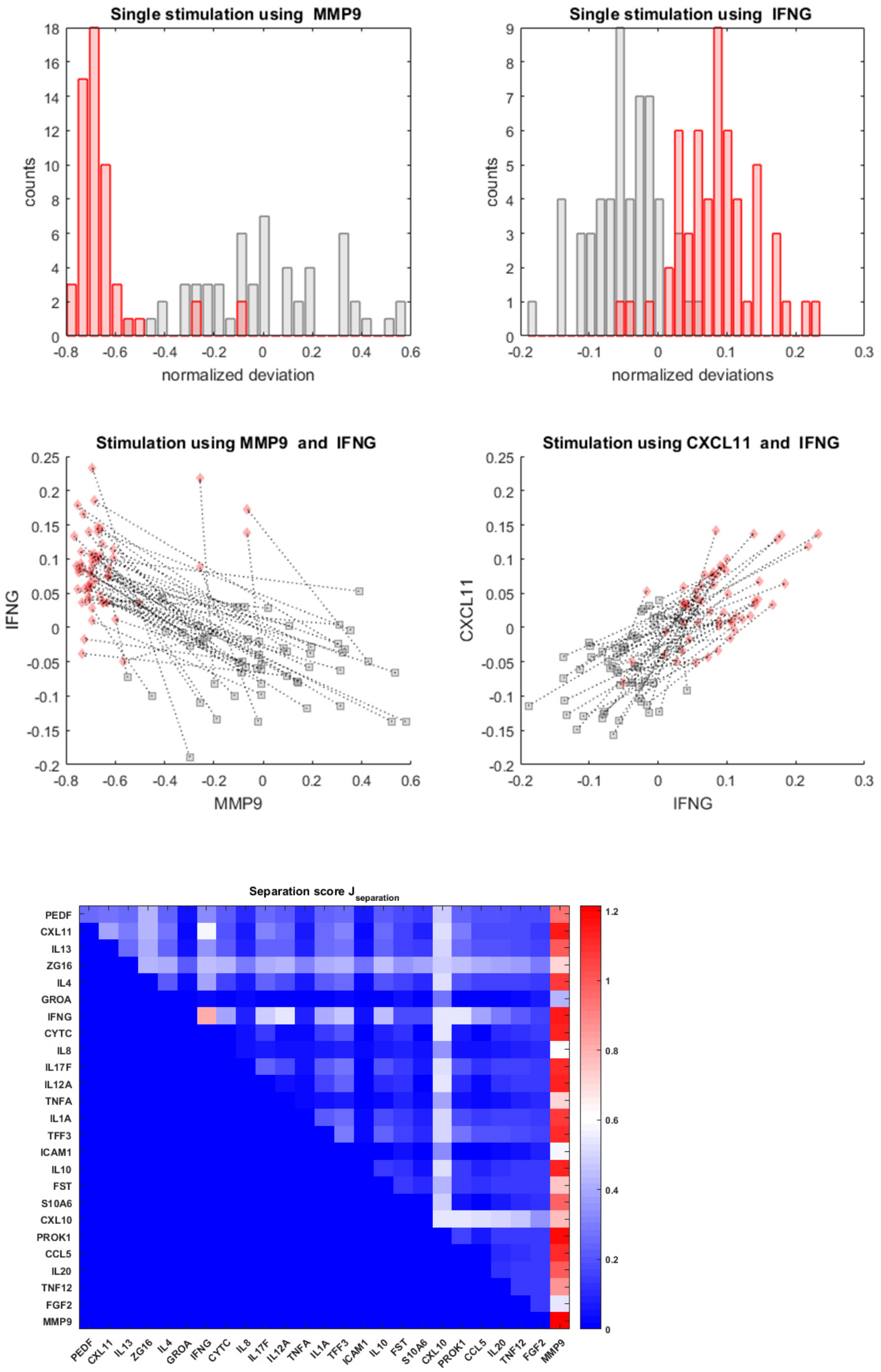


Figure S3: Finding the most discriminative proteins and protein pairs. *Top:* The top row shows histograms for the two best separating individual proteins MMP9 and IFNG having values of $J_{separation}$ 1.21 and around 0.8, respectively. The first subfigure on the second row shows the scatterplot for this pair, confirming the excellent separation offered by them together as well as individually. The second subfigure of the same row

shows that the protein pair (IFNG,CXCL11) also offers a quite good separation, although the actual separation score value is not outstandingly high (the values is close to 0.6 and thus corresponds to the white color in the heatmap in the bottom subfigure). Red diamonds stand for degrading cartilage responses, gray squares stand for healthy cartilage responses. Dotted lines between each pair of stimuli visualize how the responses are changed between stable and degrading tissue.
*Bottom:* A 25x25 dimensional heatmap illustrating a matrix with its upper part containing separation score values $J_{separation}(i,j)$ for all plausible pairs (i,j) of measured proteins. Along the diagonal the separation scores for the corresponding single proteins are shown. A detailed analysis of the matrix shows that the separation score of MMP9 alone (low right corner) is the highest, meaning that all combinations with MMP9 give a lower score value.

| **Number** | **Cytokine 1** | **Cytokine 2** | $J_{separation}$ |
|---|---|---|---|
| 1 | MMP9 | IFNG | 1,14 |
| 2 | MMP9 | CXCL11 | 1,11 |
| 3 | MMP9 | CYTC | 1,09 |
| 4 | MMP9 | PROK1 | 1,09 |
| 5 | MMP9 | IL12A | 1,08 |
| 6 | MMP9 | IL17F | 1,08 |
| 7 | MMP9 | IL10 | 1,07 |
| 8 | MMP9 | CCL5 | 1,06 |
| 9 | MMP9 | IL1A | 1,06 |
| 10 | MMP9 | TFF3 | 1,04 |
| 11 | MMP9 | IL4 | 1,02 |
| 12 | MMP9 | IL20 | 0,97 |
| 13 | MMP9 | IL13 | 0,97 |
| 14 | MMP9 | PEDF | 0,96 |
| 15 | MMP9 | S10A6 | 0,89 |
| 16 | MMP9 | CXCL10 | 0,88 |
| 17 | MMP9 | TNF12 | 0,84 |
| 18 | MMP9 | ZG16 | 0,79 |
| 19 | MMP9 | FST | 0,74 |
| 20 | MMP9 | TNFA | 0,72 |
| 21 | CXCL10 | IFNG | 0,71 |
| 22 | CXCL10 | CXCL11 | 0,70 |
| 23 | CXCL10 | IL12A | 0,69 |
| 24 | CXCL10 | IL17F | 0,69 |
| 25 | CXCL10 | IL1A | 0,68 |
| 26 | CXCL10 | IL10 | 0,68 |
| 27 | IFNG | CXCL11 | 0,68 |
| 28 | PROK1 | CXCL10 | 0,67 |

| | | | |
|---|---|---|---|
| 29 | CXCL10 | CYTC | 0,67 |
| 30 | CCL5 | CXCL10 | 0,67 |
| 31 | CXCL10 | IL4 | 0,66 |
| 32 | CXCL10 | TFF3 | 0,66 |
| 33 | CXCL10 | PEDF | 0,64 |
| 34 | IL20 | CXCL10 | 0,64 |
| 35 | CXCL10 | IL13 | 0,64 |
| 36 | CXCL10 | ZG16 | 0,62 |
| 37 | CXCL10 | S10A6 | 0,60 |
| 38 | PROK1 | IFNG | 0,60 |

Table S1: Most discriminating protein pairs sorted by the score $J_{separation}$
