## Supplementary Information 5 for "An ex vivo tissue model of cartilage degradation suggests that cartilage state can be determined from secreted key protein patterns"

### Supplementary Information 4

The results of histology, mechanical tests, GAG release and collagen II content of cartilage discs treated for 24h with DMEM* or collagenase II are presented.

### Phenotype evaluation

*Histology*

Figure S1 shows the comparison between DMEM* (untreated) and collagenase treated cartilage discs. Surface fibrillation and fissuring is observable in the collagenase treated cartilage. Loss of proteoglycan content is shown by the loss of stain intensity in the superficial zone.

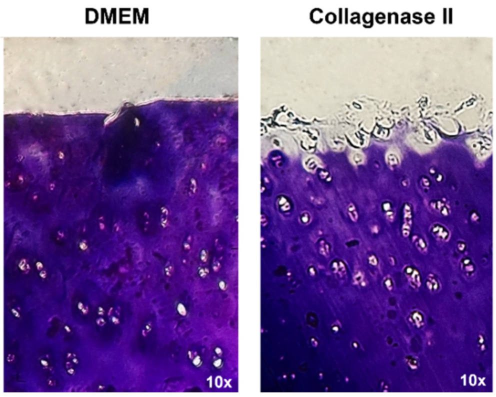


Figure S1: Representative micrographs (10x) of toluidine blue staining of DMEM* (left) and collagenase II (right) treated cartilage discs.

*Mechanical testing*

The results of the stress-relaxation experiments are shown in Figure S2. A representative stress-relaxation measurement is shown in Figure S2a. The equilibrium stresses at the end of each relaxation interval for all stable and collagenase II treated samples are depicted in Figure S2b. Collagenase treatment leads to significant reduction of the equilibrium stresses for $\varepsilon=-0.1$ ($p=0.0037$) and $\varepsilon=-0.15$ ($p<0.001$) and thus to a reduction of the tissues' aggregate elastic modulus[13].

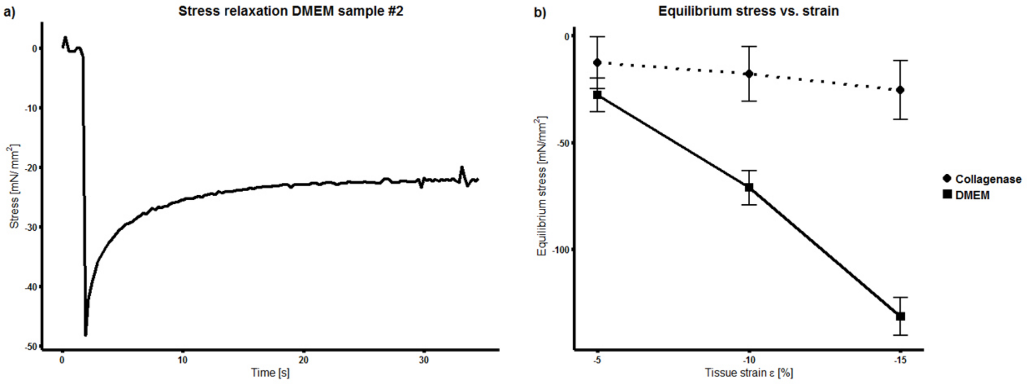


Figure S2: Results from stress-relaxation experiments of stable and collagenase II treated samples. a: A representative stress-relaxation curve after 5% compression. Equilibrium stress is the asymptotic stress at t=35 s. b: Equilibrium stress for collagenase and DMEM* treated cartilage samples (n=3) and compressive strains of 5%, 10% and 15%.

*GAG release*

The increased concentration of GAGs in the supernatant for the DMEM* and the collagenase II group is presented in Figure S3. Collagenase treatment leads to a significantly higher secretion of GAGs compared to DMEM* indicating destruction of the ECM ($p<0.001$) at 24h.

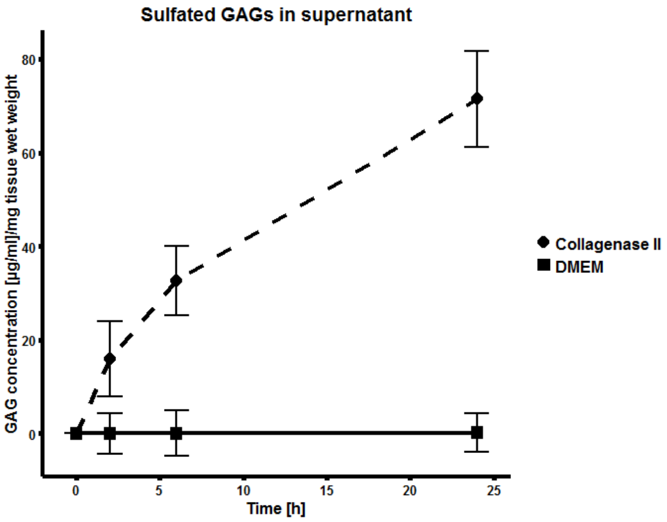


Figure S3: GAG concentration in supernatant of cartilage samples for two groups: collagenase II – dashed line with circles, DMEM* - solid line with squares. Mean values (n=3) and standard deviations are shown.

*Collagen II content*

After 24h of culture the tissue collagen content of collagenase II treated discs is significantly decreased ($p<0.001$) in comparison DMEM* treated discs, see Figure S4.

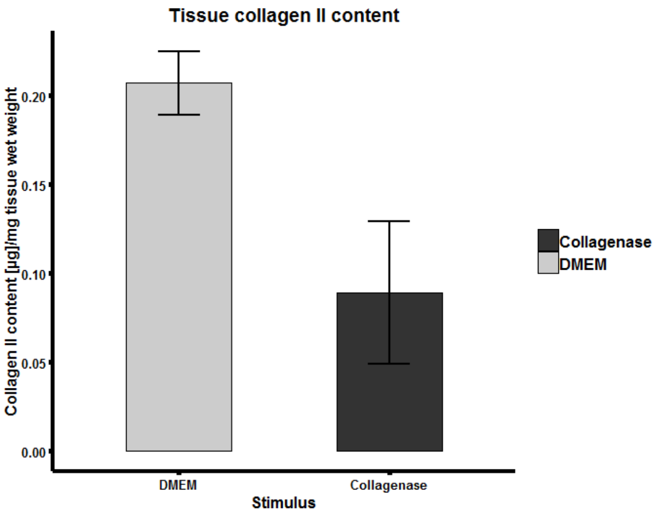


Figure S4: Tissue collagen II content per mg of tissue wet weight for two groups: collagenase II (2 mg/ml) treated discs and DMEM* treated discs. Mean values (n=3) and standard deviations are shown.
